# Title: Mechanisms of Host Cell Binding and Neurotropism of Zika Virus

**DOI:** 10.1101/350603

**Authors:** C.A. Rieder, J. Rieder, S. Sannajust, D. Goode, R. Geguchadze, R.F. Relich, D.C. Molliver, T.E. King, J. Vaughn, M. May

## Abstract

Zika virus (ZIKV) recently emerged in the Western Hemisphere with previously unrecognized or unreported clinical presentations. Here, we identify two distinct binding mechanisms of ancestral and emergent ZIKV strains featuring the envelope (E) protein residue ASN154 and viral phosphatidylserine (PS). Short (20-mer) peptides representing the region containing ASN154 from strains PRVABC59 (Puerto Rico 2015) and MR_766 (Uganda 1947) were exposed to neuronal cells and fibroblasts, expecting interactions to be representative of ZIKV E protein/cell interactions, and bound MDCK or Vero cells and primary neurons significantly above a scrambled PRVABC59 control peptide. Peptides also significantly inhibited Vero cell adsorption by ZIKV strains MR_766 and PRVABC59, indicating that we have identified a binding mechanism of ancestral African ZIKV strains and emergent Western Hemisphere strains.

Pretreatment of ZIKV MR_766 and PRVABC59 with the PS-binding protein annexin V significantly inhibited replication of PRVABC59, but not MR_766, suggesting that Western hemisphere strains are additionally utilizing PS-mediated entry to infect host cells. Taken together, these data indicate that we have identified an ancestral binding mechanism of ZIKV, and a secondary binding mechanism utilized by Western Hemisphere strains.

## Background

Zika virus (ZIKV) is a mosquito-borne *Flavivirus* that recently emerged and established endemicity in the Western Hemisphere (reviewed here^1–2^). ZIKV disease historically presented as a mild febrile illness featuring myalgia, rash, and conjunctivitis; however, novel and more severe clinical presentations and increased disease incidence were reported as the virus emerged in the South Pacific^3–4^ and the Western Hemisphere^5–7^. Since that time, case reports and animal models have implicated ZIKV in a congenital syndrome most notably featuring microcephaly^7–12^, primary encephalitis, encephalomyelitis, lyssencephaly, or Guillan-Barré syndrome^5, 13–20^, chorioamnionitis^21^, testicular infection^22^, changes in semen quality^23^, and potentially a hemorrhagic shock syndrome^24–25^. The biology and pathogenesis of ZIKV were virtually unexplored at the time of its detection in the Western Hemisphere, making rapid progress toward diagnostics, therapeutics, or vaccine development challenging in the absence of
targets^26^.

Substantial progress in the understanding of ZIKV biology has been made in a short time, and includes identification of divergent nucleotide and amino acid sites^27–30^, potential host cell ligands^31–33^, factors that impact replication kinetics^34–37^, and the development of animals models for both neurological and prenatal disease^38–39^. A recent study by Yuan *et al.* demonstrated that a single amino acid substitution within the PrM protein of Western Hemisphere strains conferred increased virulence and resulted in exacerbated pathology *in vivo*^37^. While this change confers an increased capacity for cell death and correlates with the clinical findings suggesting more severe and invasive disease, it cannot explain the newly emerged ability to directly invade the central nervous system (CNS). We sought to build upon our recent informatics analysis^27^ by utilizing the findings to identify the binding mechanism of ZIKV. We hypothesized that NAG-glycosylation of E protein ASN154 is linked to ZIKV neuroinvasiveness and that this region plays a critical role in host cell adsorption across strains.

## Results and Discussion

### Binding Motif Prediction

Structural modeling predictions of strain PRVABC59 (Puerto Rico, 2015) indicated that ASN154 is part of a linear β strand (Fig.1A). The disorder probability of this region peaks at 0.72 (Fig.1B), suggesting that this portion of the E protein is particularly dynamic and flexible. Structural and disorder probability predictions of the African type strain MR_766 (Uganda 1947) and the Nigerian strain IbH30656 (Nigeria, 1968) exhibit similar characteristics (Fig.1C, D). This region was termed the (putative) Zika virus binding motif (ZVBM).

**Figure 1.**
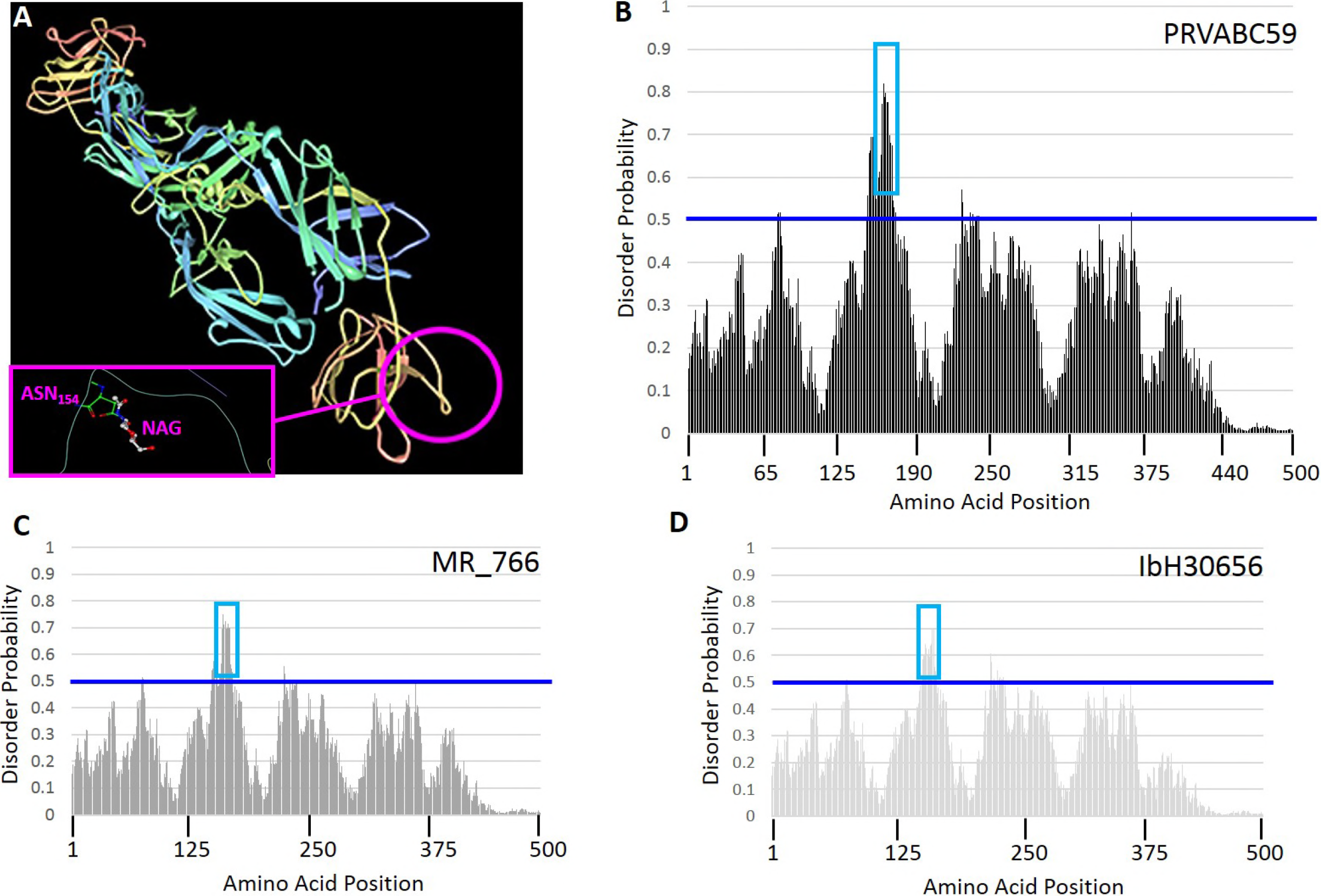
Envelope Protein Structure. (A) The predicted E protein structure indicates that the region containing ASN154 (circled) and its NAG modification (inset) is a linear β strand. Intrinsic disorder probabilities were calculated for each amino acid position in the E protein sequence from strains (B) PRVABC59, (C) MR_766, and (D) IbH30656. Probabilities above 0.5 (blue line) are considered indicative of sites representing disordered regions. The region containing ASN154 is indicated (blue box) for each strain.

#### ZVBM Binding and ZIKV Inhibition

ZVBM sequences from strains PRVABC59, MR_766, and IbH30656 were synthesized and N-terminally labelled with fluorescein isothiocyanate (FITC) (Table 1) in order to assess their capacity to bind ZIKV-susceptible and-permissive cell lines, disrupt ZIKV adsorption, and to interact with dorsal root ganglia (DRG) neurons *ex vivo*. The PRVABC59 sequence was used to generate a peptide that was modified with an NAG molecule at position 8 (equivalent to ASN154), as it natively occurs in this strain, and without carbohydrate modification. ZVBM peptides from MR_766 and PRVABC59 (NAGylated and unglycosylated) all bound Vero cells at levels significantly above those of scrambled PRVABC59 (NAGylated and unglycosylated) controls (Fig. 2A, supplemental Fig. S1), suggesting that this motif has the potential to serve as a ZIKV receptor. Binding of the MR_766 ZVBM peptides, despite a four-amino acid deletion relative to PRVABC59, suggests that the critical portion of the ZVBM is potentially contained entirely on the aminoterminus (NTD) or the carboxyterminus (CTD). A peptide representing the NTD of PRVABC59 was unable to bind Vero cells above scrambled control, whereas a peptide representing the CTD bound significantly (*P*<0.05) above the scrambled control, indicating that the putative receptor is contained entirely on the CTD of ZVBM (Fig. 2B). Pretreatment of Vero cell monolayers with the unglycosylated (″Africanized″) PRVABC59 ZVBM significantly (*P*<0.01) inhibited infectivity of ZIKV MR_766; conversely, pretreatment of Vero cells with the NAGylated (native) PRVABC59 ZVBM significantly (*P*<0.05) inhibited infectivity of ZIKV PRVABC59 (Fig. 2C, Supplemental Table S1). These findings demonstrate that adherence of ZVBM peptides to Vero cells has functional relevance, and that this motif mediates at least some host cell adsorption.

**Table 1.**
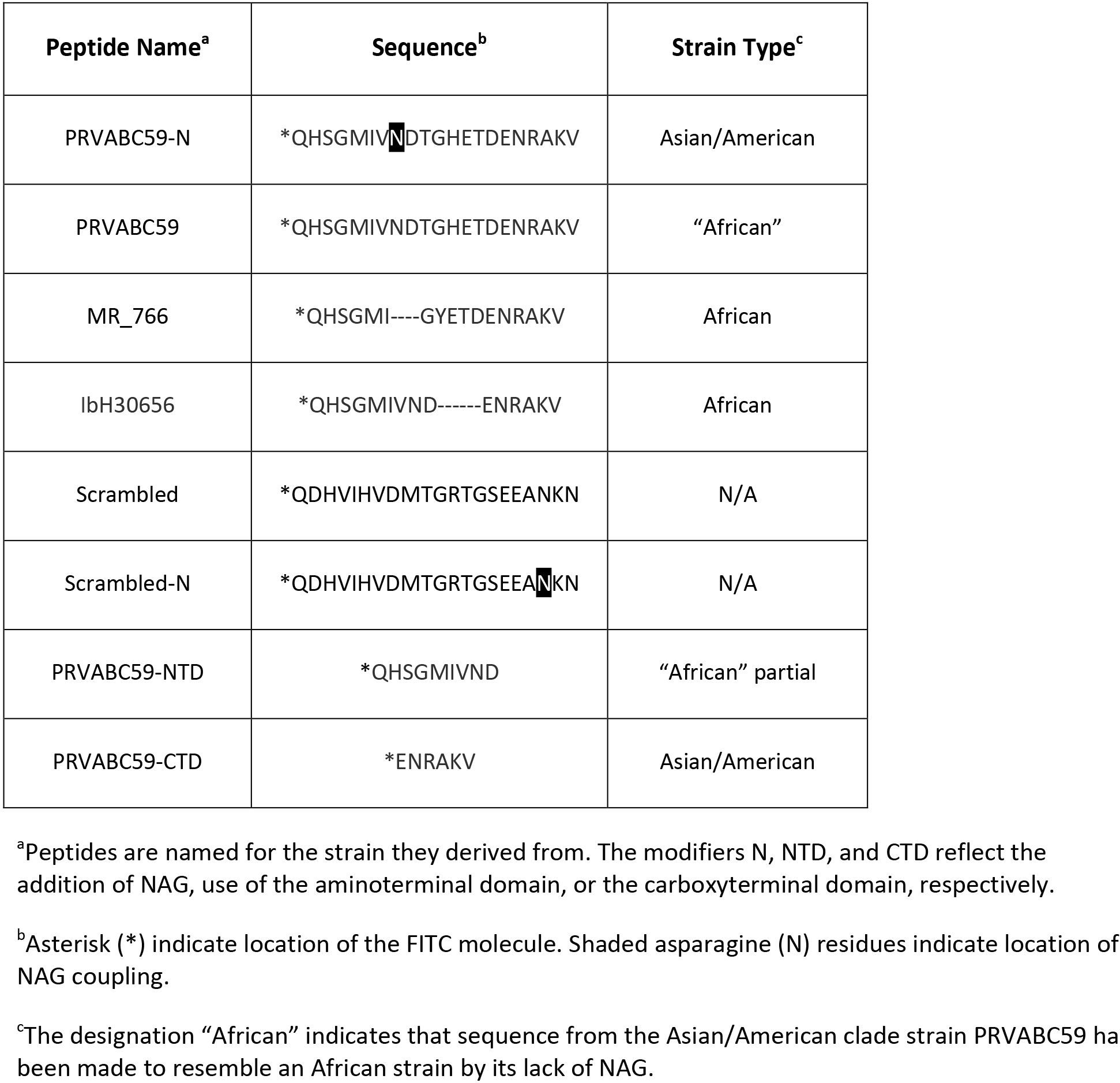
Peptide Sequences

**Figure 2.**
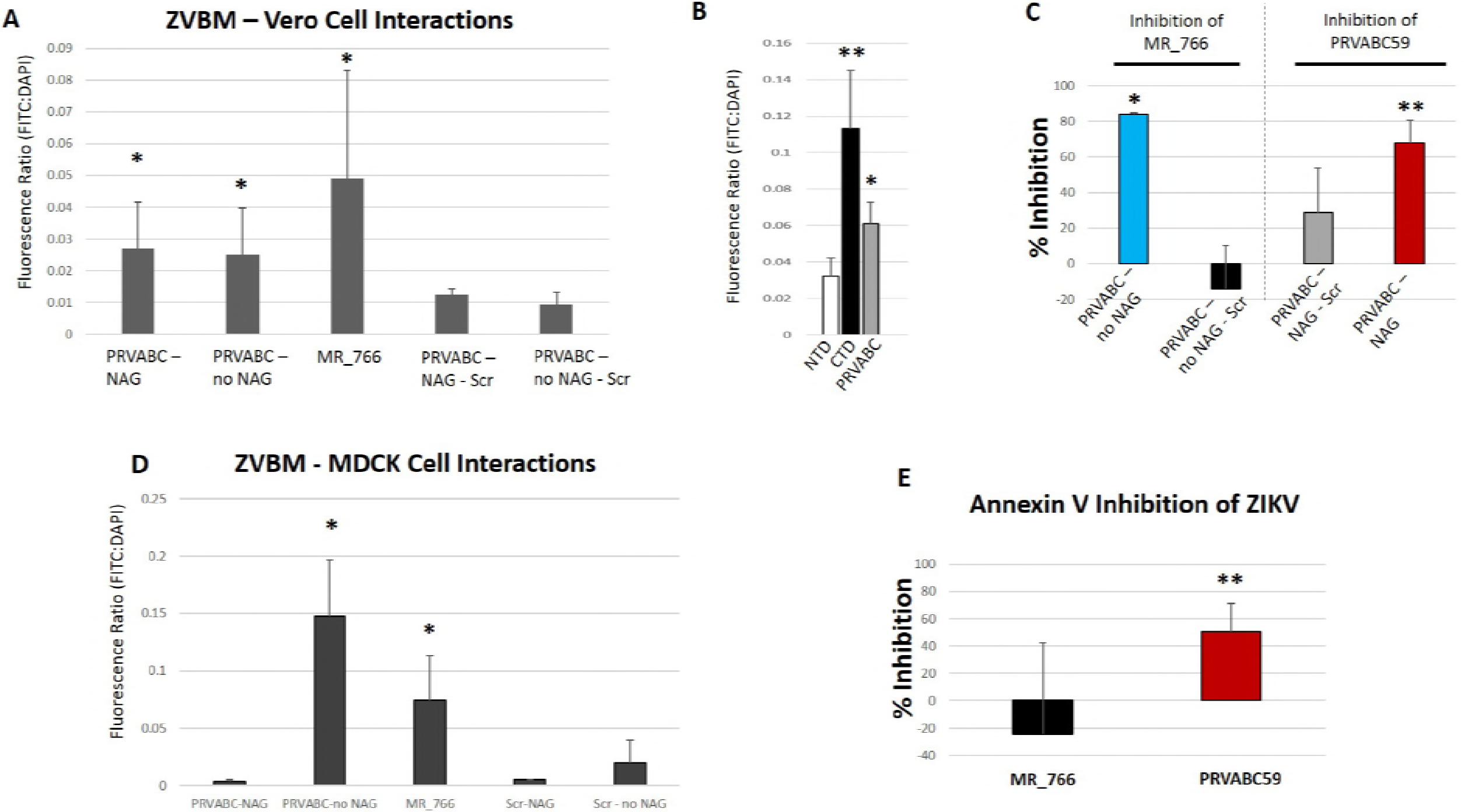
ZVBM Binding and ZIKV Inhibition. (A) NAGylated or unglycosylated ZVBM peptides from strain PRVABC59 and the type strain MR_766 bound Vero cells significantly (* *P*<0.05) above scrambled ZVBM sequences from PRVABC59, with or without NAG modification. (B) A shorter peptide representing the NTD of PRVABC59 was unable to bind Vero cells above the full-length ZVBM scrambled control; however, a peptide representing the CTD bound Vero cells at a significantly (** *P*<0.01) higher level than the NTD or scrambled control. (C) Pretreatment of Vero cells with unglycosylated ZVBM peptide (PRVABC59) significantly (* *P*<0.05) inhibited ZIKV MR_766 replication relative to untreated controls, whereas the unglycosylated scrambled control peptide was not. Similarly, pretreatment with NAGylated ZVBM peptide (PRVABC59) significantly (** *P*<0.01) inhibited ZIKV PRVABC59 replication relative to untreated controls, whereas the NAGylated scrambled control peptide was not. (D) Unglycosylated PRVABC59 and MR_766 ZVBM peptides bound MDCK cells significantly (* *P*<0.05) above unglycosylated or NAGylated scrambled controls; however, the NAGylated PRVABC59 ZVBM peptide was not. (E) Pretreatment of ZIKV strain PRVABC59 with annexin V significantly (** P<0.01) inhibited viral replication on Vero cells relative to untreated controls, whereas replication of ZIKV MR_766 following pretreatment with annexin V was unaffected.

Unglycosylated ZVBM peptides from MR_766 and PRVABC59 bound Madin-Darby Canine Kidney (MDCK) cells significantly (*P*<0.05) above (unglycosylated) scrambled control. Interestingly, NAGylated PRVABC59 did not bind MDCK cells above the NAGylated scrambled control, indicating that functionality of ZVBM as a receptor is host cell-specific (Fig. 2D). Given that MDCK cells are still permissive for ZIKV replication^40^, we hypothesized that a more generalized receptor may be contributing to viral adsorption when ZVBM is NAGylated. The association of human AXL with ZIKV adsorption^31–33^ suggests that viral phosphatidyl serine (PS) may facilitate entry into certain host cells by binding Gas6, which in turn binds AXL, as is seen with multiple viruses^41–45^. We pretreated ZIKV strains MR_766 and PRVABC59 with the PS-binding protein Annexin V prior to infection of Vero cell monolayers. Annexin V significantly (*P*<0.05) inhibited infectivity of PRVABC59 relative to untreated controls (51% reduction) but did not inhibit MR_766 (Fig.2E, Supplemental Table S1), indicating that PS-mediated ZIKV adsorption is possible for ZVBM-NAGylated (*i.e.*, Asian and American lineage) strains. While at least one additional mechanism has been described for the greater infectivity of Asian and American strains^37^, PS-mediated host cell entry is likely to contribute to this phenotype as well. The capacity of Asian and American lineage strains to utilize at least two binding mechanisms (*i.e.*, PS and ZVBM) suitably explains how AXL can serve as a host cell receptor, but animals who have undergone genetic ablation for *axl* can still serve as permissive hosts for ZIKV^46–48^.

#### ZVBM Binding to Neuronal Cells

Disease or infectivity with MR_766 following intrathecal or intracerebral inoculation *in vivo* or neuronal cell infection *in vitro* has been reported^49–53^. These findings stand in conflict with a lack of evidence for central nervous system (CNS) involvement during human disease caused by African ZIKV strains. We hypothesized that exposure to neuronal cells *ex vivo* would result in MR_766 ZVBM peptide binding, and the lack of neurological complications during Zika virus disease caused by African strains stems from an inability of these strains to penetrate into the CNS. We collected dorsal root ganglia from C57/black mice and cultured DRG neurons on coverslips. ZVBM peptides from PRVABC59 (unglycosylated), MR_766, and IbH30656 were all shown to bind 24-hour DRG neuron cultures by confocal microscopy (Figure 3A-D). Binding was not detected for the scrambled PRVABC59 control peptide. These findings were consistent with those of Annamalai *et al.*, who demonstrated a lack of neurological disease with strains lacking NAGylation at ASN154 when injected intravenously, and overt disease when the same strain was injected intracranially^54^.

**Figure 3.**
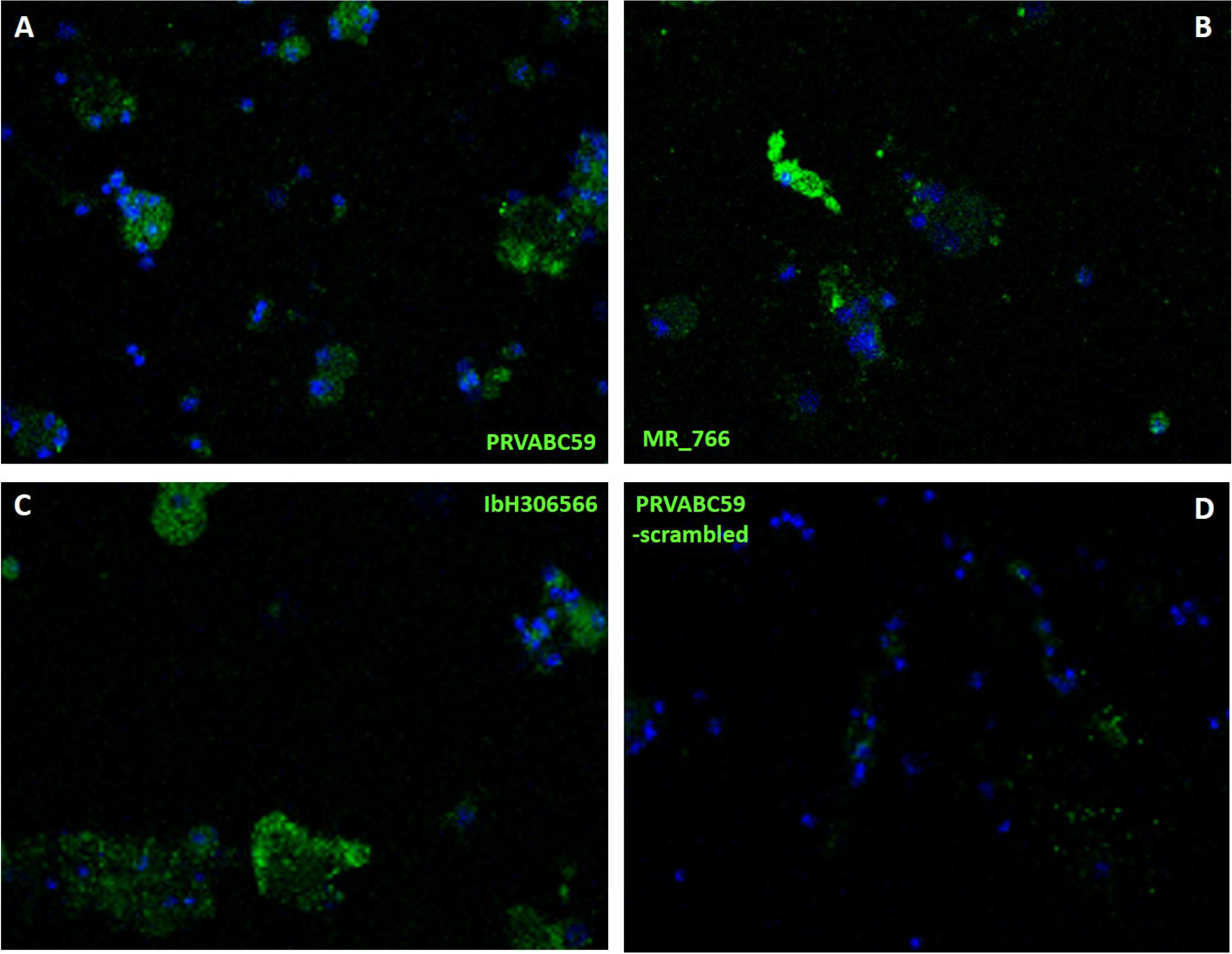
Peptide binding to DRG Neurons ***ex vivo***. Primary DRG neurons from C57 black mice (DAPI,blue fluorescence) were exposed to ZVBM peptides (FITC, green fluorescence) from (A) PRVABC59, (B)MR_766, (C) IbH30656, and (D) PRVABC59 (scrambled). Punctate green staining around the DRG nuclei was observed via laser confocal microscopy in panels A, B, and C, but was largely absent from panel D.

The outcomes of our *in vitro* and *ex vivo* studies suggest a model of ZIKV neurotropism stemming from NAGylation of the ASN154 facilitating entry into the CNS, wherein binding of certain neuronal cells occurs via the carboxyterminal portion of the ZVBM *(i.e.*, ENRAKV). This model is consistent with both the clinical disparity between ZIKV lineages and the generation of neurological disease by the African strain MR_766 when introduced directly into the CNS as previously described^37, 50^. Our findings also support previous studies that both implicate AXL as a host cell ligand for Asian/American ZIKV strains and those that show genetically ablated animals are still susceptible to infection by establishing two distinct binding mechanisms for this clade^31–33, 46–48^. These findings demonstrate the impact of NAGylation of a pathogen surface protein in the vicinity of its binding motif; namely, that there is enhanced potential to penetrate into privileged body sites. This change in posttranslational modification can therefore instantly expand the potential target tissues of infectious agents, and can be expected to similarly expand the array of clinical presentations they cause in turn.

## Methods

### Virus Isolates and Culture Conditions

African Green Monkey Kidney (Vero) cells and Madin-Darby Canine Kidney (MDCK) cells were obtained from the American Type Culture Collection. Cells were routinely propagated in Earle’s Minimum Essential Medium (EMEM) with Earle’s Balanced Salt Solution, supplemented with 10% fetal bovine serum, L-glutamine and Penicillin/Streptomycin. Cell cultures were incubated at 37°C, with 5% CO_2_ and a relative humidity (RH) of 90%. Low-passage isolates of ZIKV strains MR_766 (ATCC VR-84, Uganda) and PRVABC59 (ATCC VR-1843, Puerto Rico) were obtained from the American Type Culture Collection. Virus stocks were propagated on monolayers of Vero cells. Harvested virus lysates were clarified by low-speed centrifugation (500xg/10 min.) and stored in 1-ml aliquots at -80°C.

### Protein Analysis and Peptide Design

The Envelope protein structure was visualized using Jmol via the Protein Data Bank (PDB ID 5JHM)^55–56^, and the PDB Ligand Explorer was used to visualize the structure of N-acetyl glucosamine on ASN154. Probabilities of protein disorder at each amino acid site was estimated using PrDOS^57^. This analysis indicated that the region surrounding ASN154 constitutes a highly disordered linear epitope. Synthetic peptides representing this linear epitope including the differentially glycosylated ASN154 were generated (see Table 1) by Bachem (Bubendorf, Switzerland). The aminoterminal and carboxyterminal domains from PRVABC59 were also synthesized. Peptides were modified by the addition of an aminoterminal FITC label to allow detection and visualization.

### Primary Dorsal Root (DRG) Ganglia Neuron Culture

Adult C57/black mice were anesthetized and perfused transcardially with 4°C 1× PBS. Cervical, thoracic and lumbar DRGs were dissected in Ca^++^/Mg^++^-free Hank’s basic salt solution (HBSS) and dissociated as previously described^58^. DRGs were cultivated on laminin/polyD-lysine coated EZ slides (MilliporeSigma, Burlington, MA) for 18-24 hours in F-12 medium (Gibco, ThermoFisher Scientific, Waltham, MA) supplemented with 10 % fetal bovine serum, 1 % penicillin/streptomycin at 37° C/5% CO_2_. Bound peptides were visualized with a Leica TCS SP5 confocal laser scanning microscope.

### Peptide Binding Assays

Vero cells and MDCK cells, both of which are permissive for all Zika strains, were grown to 80% confluency in 48-well plates. Following the removal of medium, wells were blocked with 10% fetal bovine serum for 30 minutes at 37° C. Peptides (100 μg/mL) were incubated with Vero or MDCK cells for 1 hour at 37° C. Unbound peptides were removed by washing with 1× PBS, and mammalian cells were counterstained with DAPI (diluted 1:300) to control for minor variations in monolayer populations. Bound peptides (FITC) were detected at 485/490 (excitation/emission), and cells were quantified at 350/460. Data are presented as FITC:DAPI ratios. Statistical significance was measured by analyses of variance, and by Fisher’s Protected Least Significant Difference test for posthoc comparisons when main effects were significant (GraphPad Prism v. 6.0). Primary DRG neurons grown on coverslips were incubated with 100 μg peptide for 1 hour at 37° C. Unbound peptides were removed by washing with HBSS, and DRG neurons were counterstained with DAPI (diluted 1:300). Bound peptides were visualized using Keyence BZX-700 inverted widefield digital microscope.

### Viral Adsorption Inhibition Assays by ZVBM Peptides

Vero cells were propagated in 48-well plates (seed concentration-1e5 cells/well) for 24-hr. Resulting monolayers (85% confluence) were washed twice with warmed PBS and incubated for 2 hours (37° C, 5% CO_2_, 90% RH) with 0.1 ml volumes of either PBS (Negative Control), or PBS containing 467 ug of the selected ZVBM peptide. Following treatment, PBS or peptide was decanted, and monolayers washed twice with warmed PBS. ZIKV stocks (stock concentrations: Strain MR_766 – 3.16 e7 TCID_50_/ml; Strain PRVABC59 – 2e7 TCID_50_/ml) were serially diluted in serum-free Dulbecco’s Minimum Essential Medium (DMEM). Host cell monolayers in treated, or untreated plates were inoculated with either strain MR_766 or PRVABC59 (0.1 ml/well, 5 wells per dilution, N = 3 replicates each) and incubated for 2 hr. After inocula were removed, wells were supplemented with 0.5 ml EMEM growth medium and returned to the incubator. Virus cytopathogenic effects (CPE) were monitored and scored over a period of 10-12 days and the resulting virus titers calculated as TCID_50_/ml. Statistical significance of changes in virus titer as a result of peptide pretreatment versus untreated control was measured by *Student’s* T-test (GraphPad Prism v. 6.0).

### Virus Treatment with Annexin V

Annexin V (AbCam, Cambridge, MA) was dissolved in PBS (2335 ug/ml) and filter-sterilized (0.2 um). ZIKV stocks MR_766 and PRVABC59 were then serially diluted in either PBS, or PBS-Annexin V. Dilutions were incubated for 2 hours (37° C, 5% CO_2_, 90% RH). Host cell monolayers, prepared in 48-well plates as previously described, were inoculated with respective virus dilutions (0.1 ml/well, 5 wells per dilution, N = 3 replicates each). Virus CPE were scored over a period of 10-12 days and the resulting virus titers calculated as TCID_50_/ml. Statistical significance of changes in virus titer as a result of Annexin V pretreatment versus untreated control was measured by *student’s* T test (GraphPad Prism v. 6.0) for each ZIKV strain.

### Ethical Assurance Statements

All methods were carried out in accordance with relevant guidelines and regulations. Collection of DRG neurons was performed in accordance with a protocol approved by the University of New England’s Institutional Animal Care and Use Committee.

## Acknowledgements

This work was supported by intramural awards from the University of New England Center of Excellence for Neuroscience (CEN) an the Office of the Vice President for Research and Scholarship. The authors wish to thank Peter Caradonna (CEN Histology and Imaging Core), Denise Giuvelis (CEN Behavioral Core), and Joshua Havelin for their assistance.

## Author Contributions

Peptide motif design and experimentation (CAR, JR, SS); structural and informatics analysis (MM, RFR); neuronal cell extraction development, binding experimentation, confocal imaging parameters (DG, RG, DCM, CAR, SS); viral replication inhibition studies (JV, CAR); Study design, management, execution, and data analysis (MM, RFR, TEK); manuscript preparation (CAR, MM)

**Additional Information:**

## Supplemental Material

**Supplemental Table S1.**
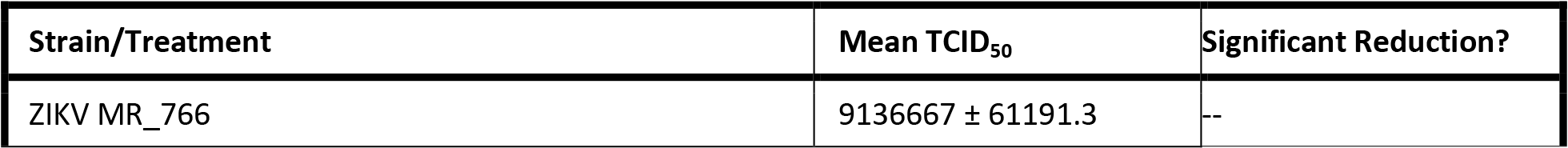
ZIKV Adsorption Inhibition

**Supplemental Table S1a.**
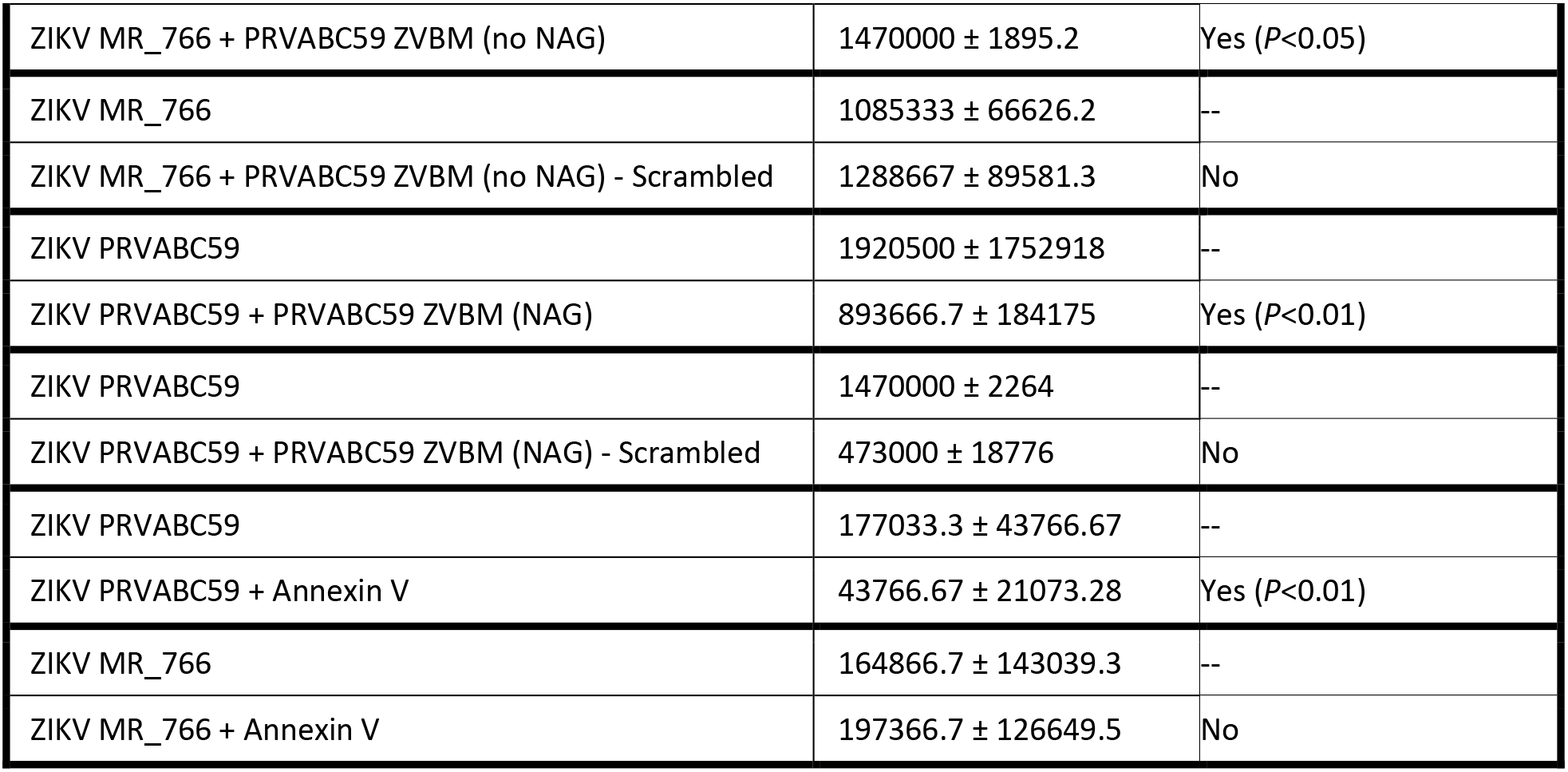

**Supplemental Figure S1.**
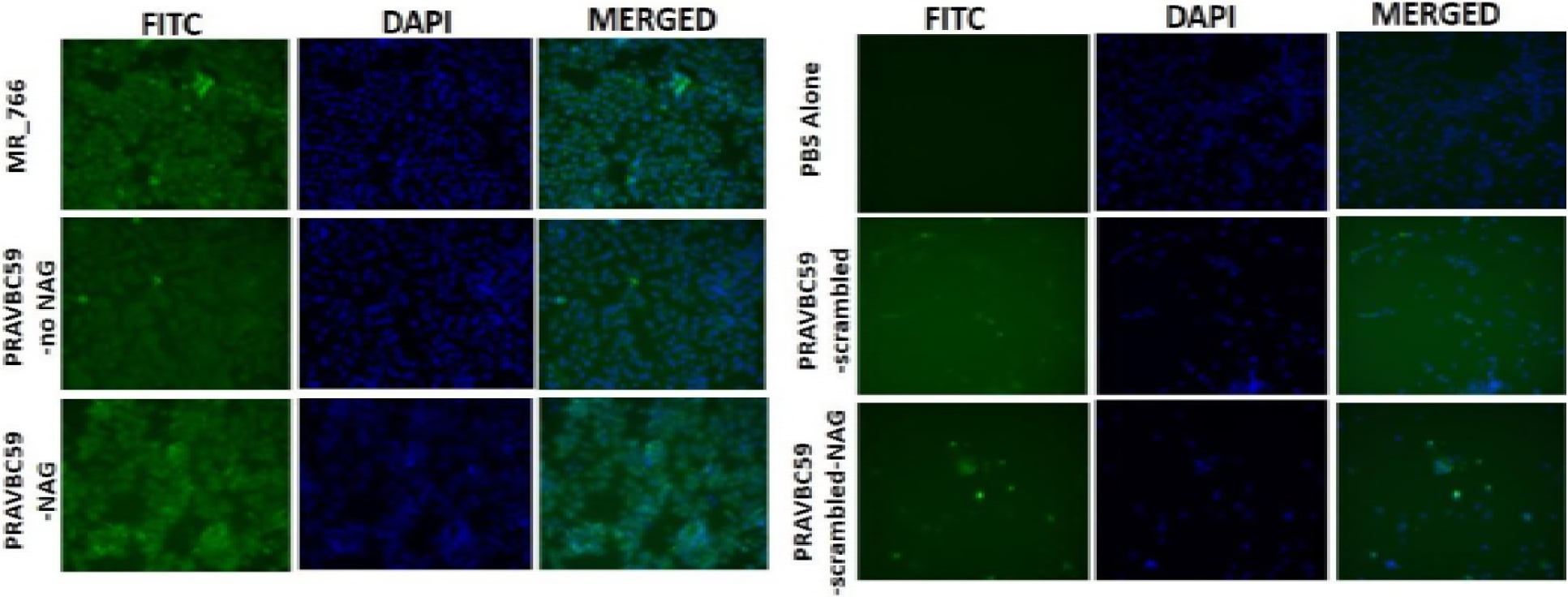
Peptide binding to Vero Cells. Vero cell monolayers (85% confluency) (DAPI,blue fluorescence) were exposed to ZVBM peptides (FITC, green fluorescence) from PRVABC59 (NAGylated and unglycosylated), MR_766, and scrambled PRVABC59 peptides (NAGylated and unglycosylated). Co-localization of FITC and DAPI staining was observed for glycosylated and unglycosylated PRVABC59 and MR_766 peptides, but was largely absent from scrambled control peptides.

